# Melanin regulates mitochondrial dynamics, metabolism and inflammatory signaling to protect the retina

**DOI:** 10.64898/2026.05.15.724948

**Authors:** Md Jahirul Islam, Yong-Su Kwon, Julie Munsoor, Christopher Wu, Lucy Wang, Min Zheng, Zongchao Han

## Abstract

Albino individuals are clinically recognized to exhibit heightened susceptibility to light-induced retinal injury, yet the cellular and metabolic mechanisms underlying this vulnerability remain poorly defined. Here, we investigated whether retinal pigment epithelium (RPE) pigmentation governs mitochondrial structure, metabolism, and inflammatory responses that ultimately determine retinal resilience to blue light stress. Using pigmented (C57BL/6J) and albino (Balb/c) mice, we demonstrate that albino animals exhibit markedly increased retinal phototoxicity following blue light exposure, manifested by fundus lesions, outer nuclear layer (ONL) disruption, and structural degeneration evident by OCT. Primary RPE cultures derived from albino mice exhibited profound difference in mitochondrial morphology, characterized by increased mitochondrial number, reduced size, and enhanced fragmentation, accompanied by elevated mitochondrial DNA copy number. These structural changes correlated with transcriptional skewing toward mitochondrial fission (increased *Drp1*) and suppression of mitochondrial fusion (*Mfn1, Mfn2, OPA1*). Functionally, albino and depigmented RPE displayed impaired oxidative phosphorylation, reduced ATP production, and diminished reliance on mitochondrial pyruvate carrier (MPC)–dependent metabolism. In parallel, albino RPE demonstrated cell-cycle accumulation at G2/M and heightened basal and blue light-induced secretion of pro-inflammatory cytokines, particularly IFN-β1, IL-6, and TNF-α. Importantly, exogenous melanin supplementation partially restored mitochondrial fusion gene expression, pyruvate-dependent respiration, and inflammatory restraint. Together, these findings identify melanin as a critical regulator of RPE mitochondrial architecture, metabolic substrate utilization, and inflammatory signaling, providing a mechanistic framework to explain enhanced photo-vulnerability in the albino retina. These insights establish pigmentation-dependent mitochondrial metabolism as a determinant of retinal resilience and suggest mitochondrial bioenergetics as a therapeutic target.

## Introduction

The retinal pigment epithelium (RPE) is a multifunctional specialized epithelial monolayer that performs essential functions required for retinal homeostasis and visual function[1], including photoreceptor outer segment phagocytosis[2], recycling of visual chromophores[3], regulation of ion and metabolite transport[4], maintenance of the outer blood–retinal barrier, and modulation of innate immune responses in the outer retina[5, 6]. Due to its intimate anatomical and metabolic coupling with photoreceptors, the RPE is continuously exposed to high levels of oxidative stress generated by light exposure, mitochondrial respiration, and lipid-rich phagocytic activity[7, 8]. Failure of RPE homeostasis is therefore a central event in many retinal degenerative conditions, ultimately leading to photoreceptor dysfunction and vision loss[9, 10].

A defining feature of the RPE is its high content of melanosomes, which accumulate during development and undergo age-related changes, including a gradual decline in melanin content and increased formation of melanolipofuscin[11] [12, 13]. RPE melanin, the dominant constituent in the melanosome, serves as a broad-spectrum absorber of visible and ultraviolet light and functions as a potent scavenger of reactive oxygen and nitrogen species, thereby limiting photo-oxidative damage[14-16]. In addition to its light-screening properties, melanin can bind redox-active metals, buffer free radicals, and modulate local oxidative chemistry within the RPE cytoplasm[17]. Clinically and experimentally, loss or absence of melanin as observed in albinism, normal aging[18], and retinal degenerative diseases such as age-related macular degeneration (AMD) is strongly associated with increased retinal vulnerability to light exposure, heightened oxidative stress, and progressive photoreceptor degeneration[13, 19, 20]. Despite these associations, the intracellular mechanisms by which melanin deficiency alter RPE physiology remain incompletely understood.

Blue light (400–500 nm) represents a particularly damaging component of the visible spectrum due to its high photon energy and strong capacity to drive photo-oxidative reactions within photoreceptors and RPE mitochondria[21-23]. Blue light exposure induces mitochondrial reactive oxygen species (ROS) generation, lipid peroxidation, DNA damage, and inflammatory signaling in retinal cells[24-26]. Epidemiological data and experimental models consistently demonstrate that albino animals and individuals exhibit markedly increased sensitivity to blue light–induced retinal damage, characterized by accelerated photoreceptor loss, outer retinal inflammation, and impaired visual function[27, 28]. While reduced optical filtering by melanin is widely considered a primary contributor, increasing evidence suggests that melanin may exert active regulatory roles that extend beyond passive photoprotection[29, 30].

Mitochondria are central to RPE physiology, supplying the substantial ATP demands required for ion transport, phagocytosis of photoreceptor outer segments, visual cycle enzymatic reactions, and metabolic support of photoreceptors[31, 32]. In addition to energy production, mitochondria act as key signaling hubs that integrate redox balance, metabolic status, and inflammatory responses[33]. Mitochondrial dysfunction manifested by impaired oxidative phosphorylation, altered substrate utilization, excessive ROS production, and disrupted mitochondrial dynamics has emerged as a unifying pathogenic mechanism in retinal degenerative diseases and light-induced retinal injury[34, 35]. Excessive mitochondrial fission driven by dynamin-related protein 1 (Drp1) promotes mitochondrial fragmentation, bioenergetic inefficiency, and activation of stress-associated inflammatory pathways, whereas mitochondrial fusion mediated by mitofusins (Mfn1 and Mfn2) and optic atrophy 1 (OPA1) supports efficient oxidative metabolism and cellular stress tolerance[36-38].

Recent ultrastructural and biochemical studies suggest that melanin and melanosomes may physically and functionally interact with mitochondria within RPE cells, influencing localized redox environments and metabolic activity[39-41]. Furthermore, emerging work highlights the importance of mitochondrial substrate selection in RPE health, particularly the role of mitochondrial pyruvate carrier (MPC)–mediated pyruvate imports in sustaining oxidative phosphorylation and retinal metabolic homeostasis[42-44]. Disruption of MPC function compromises ATP production and predisposes retinal cells to oxidative damage, yet whether pigmentation status modulates mitochondrial substrate utilization remains unknown. In addition to metabolic dysregulation, mitochondrial stress is increasingly recognized as a trigger for inflammatory signaling in the RPE. Mitochondrial dysfunction can promote type I interferon responses, cytokine secretion, and breakdown of immune privilege, processes that contribute to retinal degeneration[9, 10, 45]. Whether melanin deficiency amplifies mitochondrial stress-induced inflammatory responses in RPE has not been systematically investigated.

Here, we test the hypothesis that pigmentation status fundamentally reprograms RPE mitochondrial architecture, metabolic substrate utilization, and inflammatory signaling, thereby dictating retinal sensitivity to blue light–induced injury. Using complementary in vivo retinal phototoxicity models, ex vivo RPE analyses, and Seahorse XF metabolic flux measurements across pigmented, depigmented, albino, and re-pigmented systems, we identify a melanin-dependent regulatory axis linking mitochondrial dynamics, pyruvate-driven oxidative metabolism, and inflammatory restraint to retinal protection.

## Methods

### Induction of Retinal Photodamage by Blue Light Exposure

C57BL6/J (B6) and Balb/c mice were purchased from the Jackson Laboratory. All experiments were conducted in accordance with the policies of the Institutional Animal Care and Use Committee (IACUC) at The University of North Carolina at Chapel Hill (UNC-CH) and were maintained following the Association for Research in Vision and Ophthalmology (ARVO) statements regarding the use of animals in ophthalmic and vision research. All mice in experiments were 6 − 8 weeks old postnatal. Before blue light exposure mice were dark-adapted for 16 h. Next, the pupils of the mice were dilated with 1% tropicamide (Bausch & Lomb Inc., Tampa, FL, USA) under dim red light. Acute retinal damage was induced by exposing mice to 900 to 10,000 lx of diffuse blue LEDs lights (Super Bright LEDs, 78 lm/W, 64 W, 5050 SMD LEDs) using blue light exposure system for 3 h per day over 3 consecutive days. The mouse eyes were imaged using an imaging system of Micron III fundoscopy system (Phoenix Research Laboratories, Pleasanton, CA, USA) and the position was adjusted until a clear image of the fundus filled the screen and the OCT was visualized. Ketamine (85mg/kg)/Xylazine (10mg/kg) was used for anesthesia in all cases before imaging. The mice were placed on a warm bed until fully awake and kept in a temperature-controlled dark room.

### Mouse Primary RPE Cells Culture and Isolation of RPE Melanin

Primary RPE (pRPE) cells were obtained from 8 – 10-weeks old B6 and Balb/c mice eyeball after euthanasia was performed using isoflurane exposure and decapitation. pRPE cell isolation was conducted using the following procedure. Briefly, the extracted eye globes were rinsed with 70% ethanol and washed with PBS solutions. The cornea, lens, and retina were removed, and the posterior eyecup was incubated with trypsin-EDTA (0.25%, Gibco) at 37 °C for 20 min. Subsequently, the pRPE cells were harvested by gently scraping off Bruch’s membrane. The collected pRPE cells were rinsed twice with DMEM (Gibco) containing 10% (vol/vol) fetal boine serum (FBS) (Gibco) and incubated in a 24 well cell culture dish with DMEM containing 10% FBS and 1% of the penicillin streptomycin stock solution (10,000 units/ml penicillin, 10mg/ml streptomycin in 0.85% saline; P4333, Sigma Aldrich, UK) at 37 °C under 95% humidity and 5% CO2. After two days of cell culture, residual unattached pRPE cells and debris were gently removed with a PBS buffer solution, and then the cell culture medium DMEM was changed every 5 days.

To collect melanin from B6 and Balb/c mice RPE cells, the isolation method was followed with slight modifications as described [48]. Briefly, collected pRPE cells were suspended in a hypotonic buffer containing protease inhibitor cocktail (MilliporeSigma Calbiochem, 53913110VL). The pRPE cells underwent disruption with a Dounce tissue homogenizer. The whole cell lysate was then centrifuged at 3,000 g for 5 min and re-suspended in a buffer solution containing 50 mm Tris-HCl and 150 mm KCl (pH 7.4). Subsequently, the cell lysate was layered on top of a discontinuous OptiPrep gradient (50%, 35%, 30%, 20%, and 15%) and centrifuged at 135,000 g for 1 h at 4 °C (Beckman L8-80 m Ultracentrifuge, USA). For fractionation analysis each of five layers (Supplementary figure) was collected from top and sent for EPR analysis to our collaborator at NC state university core facility at 4°C temperature. The melanin pellet was recovered, purified with deionized water three times by centrifugation (Thermo Scientific Sorvall ST 16R centrifuge, USA) at 13,000 rpm for 15 min. The RPE melanin was finally redispersed in deionized water and stored at − 20°C temperature fridge.

### Whole cell imaging

pRPE cells were cultured or seeded on 12-mm microscope cover glasses (Deckglaser) in 24-well plates. Once cells are grown, passaged and ready for imaging cells were first rinsed in 1xHBSS, fixed in ice-cold PBS containing 4% formaldehyde for 30 minutes at 4°C and washed three times in 1x PBS. Blocking and permeabilization were achieved by incubation with 5% normal goat serum (NGS; G9023, Sigma Aldrich, UK) in 1x Phosphate Buffered Saline with Triton X-100 (PBST, Sigma Aldrich, UK) for one hour prior to the addition of primary antibody (prepared in blocking buffer) which was incubated overnight at 4°C. The following primary antibodies were used: ZO-1 (1:100, RRID: AB_2533456, Thermo Fisher Scientific, UK), RPE65 (1:100, RRID: AB_2181006, Abcam, UK). The following day, cells were washed in 0.05% PBST and incubated with the Alexa Flour 532-labeled goat anti-mouse IgG (Invitrogen) at a dilution of 1:200 for 1 hour. After rinsing the secondary polyclonal antibodies, cover glasses were placed on DAPI mounting solution (Vector laboratories, Burlingame, CA, USA) on glass slides. Fluorescence imaging was performed using a Confocal Microscope.

Isolated RPE tissue and cultured primary RPE cells were used for TEM preparation and imaging. Briefly, tissue/cell suspension were fixed for 1 hour at room temperature in 4% paraformaldehyde (PFA) in 0.15M sodium phosphate buffer pH 7.4. Samples were then washed 3 × 15 minutes (min) in 0.1M cacodylate buffer, stained in 1% OsO4 for 1 hour, followed by 3 × 15 min washes in ddiH20 and one wash in 50% ethanol for 5 min. Then samples were en-bloc stained with 2% uranyl acetate for 30 min. Successive dehydration washes were performed in 50, 70 and 95% ethanol 2x each for 10 min. Samples were then transferred to Wheaton glass jars where propylene oxide washes were performed 2 × 15 min each. A solution of 50/50 propylene oxide/EPON was applied overnight. The following day 25/75 propylene oxide/EPON was applied for 2 hours. Then 100% EPON was applied for 2 hours and fresh 100% EPON was applied for at least 24 hours and allowed to completely polymerize at 60°C. Blocks were trimmed to the tissue and 500µm sections were cut using a Leica EM UC7 (Leica Microsystems) and a Diatome diamond knife (Electron Microscopy Sciences). Sections were mounted on glass slides, stained with 1% toluidine blue O in 1% sodium borate. Ultrathin sections (80nm) were cut and mounted on 300 mesh copper grids. The grids were stained with 2% ethanolic uranyl acetate for 8 min, followed by staining with lead citrate for 5 min to add additional contrast. Tissue samples were visualized using a Philips Tecnai 12 at 120 kV and images were taken using a Gatan Rio 16 CMOS camera with Gatan Microscopy Suite software. Sample prep and imaging was performed at the UNC Hooker Imaging Core.

### Seahorse extracellular flux analysis of mitochondrial respiration

Using a Seahorse extracellular flux analyzer, we monitored real-time oxygen consumption rates (OCR) and extracellular acidification rates (ECAR) in RPE cells. After a 48-h incubation either with growth medium or growth medium containing mitochondrial energy transfer inhibitors: UK5099 (2 uM), Etomoxir (4 uM), or BPTES (3 uM) from corresponding well were removed, leaving 50 µL of media. And cells were washed twice with pre-warmed assay medium. Cells were incubated in 37°C incubator without CO_2_ for 1 h to allow to pre-equilibrate with the assay medium. Load pre-warmed oligomycin, FCCP, rotenone & antimycin A into injector ports A, B and C of sensor cartridge, respectively. The final concentrations of injections were 1.5 µM oligomycin, 1.5 µM FCCP, and 0.5 µM rotenone & antimycin A. The machine was calibrated and the assay was performed using glycolytic stress test assay protocol as suggested by the manufacturer (Seahorse Bioscience, Billerica, MA, USA). OCR and ECAR were detected under basal conditions followed by the sequential addition of oligomycin, FCCP, as well as rotenone & antimycin A. This allowed for an estimation of the contribution of individual parameters for basal respiration, proton leak, maximal respiration, spare respiratory capacity, non-mitochondrial respiration and ATP production.

Freshly isolated RPE tissue were incubated for 2h with growth medium before changing to assay medium for OCR analysis.

Live imaging was performed on a Zeiss LSM 980 confocal microscope equipped with a temperature- and CO2-controlled incubation chamber and a 20X (0.8 NA) objective lens. Images were acquired at baseline and after a 10 min incubation with 5 µM Antimycin A.

### Gene expression and mtDNA copy number

Total RNA was isolated from RPE tissue and primary RPE cells using an RNeasy Plus Mini kit (QIAGEN, Hilden, Germany). cDNA was prepared from 1 µg of total RNA. Thereafter, real-time quantitative PCR (qPCR) was performed using SYBR Green Fast mix (Applied Biosystems) and primers specific to target genes provided below in the StepOnePlus™ Real-Time PCR system (Applied Biosystems). The expression levels of target mRNAs were analyzed using the ddCt method and were normalized to the expression levels of mouse GAPDH, a housekeeping gene used as an endogenous control.

Drp1 Fw: TCAGATCGTCGTAGTGGGAA

Drp1 Rw: TCTTCTGGTGAAACGTGGAC

Mfn1 Fw: GCTGTCAGAGCCCATCTTTC

Mfn1 Rw: CAGCCCACTGTTTTCCAAAT

Mfn2 Fw: ATGTTACCACGGAGCTGGAC

Mfn2 Rw: AACTGCTTCTCCGTCTGCAT

Opa1 Fw: ATACTGGGATCTGCTGTTGG

Opa1 Rw: AAGTCAGGCACAATCCACT

Total DNA was isolated from RPE tissue and primary RPE cells using Puregene Cell and Tissue Kit (Qiagen) and was amplified using specific primers for mtCOXII and 18S by real-time PCR using the Power SYBR Green RT-PCR kit (Applied Biosystems). The mtDNA copy number was calculated using 18S amplification as a reference for nuclear DNA content. Real-time PCR was performed with the following primers:

18S Fw: CATTCGAACGTCTGCCCTATCA

18S Rw: GGGTCGGGAGTGGGTAATTTG

COXII Fw: GCCGACTAAATCAAGCAACA

COXII Rw: CAATGGGCATAAAGCTATGG

### Cell cycle analysis

Ice-cold 95% ethanol supplemented with 0.5% Tween-20 was added to primary RPE cell suspensions to a final concentration of 70% ethanol. The fixed cells were pelleted and washed in 1% BSA–PBS solution. The cells were resuspended in 1 ml of PBS containing 11 Kunitz U/ml RNase and were incubated at 4 °C for 30 min, washed once with BSA–PBS, and resuspended in a 50 μg/ml PI solution. The cells were incubated at 4 °C for 30 min in the dark and were then filtered through 35 mm mesh, and the DNA content was determined using a BD LSR Fortessa flow cytometer (BD Bioscience) within 1 h of staining. The cellular DNA content was analyzed using the FlowJo v9 software.

### ERG of mouse eyes

Pigmented and albino mice were dark-adapted overnight before ERG recording and anesthetized by intraperitoneal injection of ketamine (85 mg/kg) and xylazine (10 mg/kg) (Covertrus). Pupils were dilated with 1% tropicamide ophthalmic solution (Henry Schein) under dim red illumination. GenTeal lubricant gel (Alcon) was applied to ensure adequate electrical contact and to prevent corneal dehydration prior to electrode placement[46]. Long-duration ERG recordings were performed using a Celeris high-throughput electrophysiology system (Diagnosys LLC, USA) to assess the RPE-derived c-wave response at a flash stimulus intensity of 0.9 cd·s/m^2^.

### Cytokine ELISA

The inflammatory effects of blue light on primary RPE cells were assessed by quantifying the production of inflammatory cytokines, IFN-β1, IL-1 β, TNF-*α* and IL-6, utilizing commercially available ELISA kits, specifically R&D Systems Inc., Minneapolis, MN, USA) following the manufacturer’s instructions.

### Statistical Analysis

The results from all investigations were expressed as the mean ± standard error of the mean (SEM) derived from a minimum of three independent experiments. Statistical and kinetic analyses were conducted using GraphPad Prism 9.0 software (La Jolla, CA, USA). Multiple group comparisons were assessed through one- and two-way ANOVA as deemed appropriate, with statistical significance established at a p-value below 0.05 (P < 0.05).

## Results

### Albino (Balb/c) mice exhibit increased susceptibility to blue light–induced retinal damage

To compare retinal vulnerability to blue light exposure between pigmented and albino strains, pigmented (B6) and Albino (Balb/c) mice were subjected to a controlled phototoxicity paradigm (460 nm; 900–10,000 lux; 3 h/day for 3 consecutive days). Fundus imaging and OCT assessment were performed on day 5 following the final exposure. Bright-field (BF) fundus photography, fluorescein angiography (FA), and OCT images were analyzed across different exposure between Albino and pigmented mice. Albino mice demonstrated significantly greater light-induced retinal damage than their pigmented counterparts (Fig 1A). In albino animals, outer nuclear layer (ONL) discrete white fundus lesions, indicative of photoreceptor/outer retinal injury, became apparent at exposure intensities as low as 2000 lux. Correspondingly, OCT cross-sections revealed focal disruptions and thinning of the beginning at the same intensity (Fig 1B). The severity of photodamage increased progressively with higher irradiance levels (4000–10,000 lux), characterized by broader areas of hyperreflectivity and ONL disorganization. In contrast, pigmented mice displayed substantial resistance to blue light stress at intensities up to 2000 lux, exhibiting minimal to no white fundus spots and preserved ONL structure on OCT. Only at higher light intensities pigmented retinas begin to show mild photodamage, and even then, the extent remained markedly reduced compared with albino mice. Together, these results demonstrate that albino Balb/c mice are significantly more susceptible to blue light–induced retinal injury than pigmented B6 mice.

**Figure 1.**
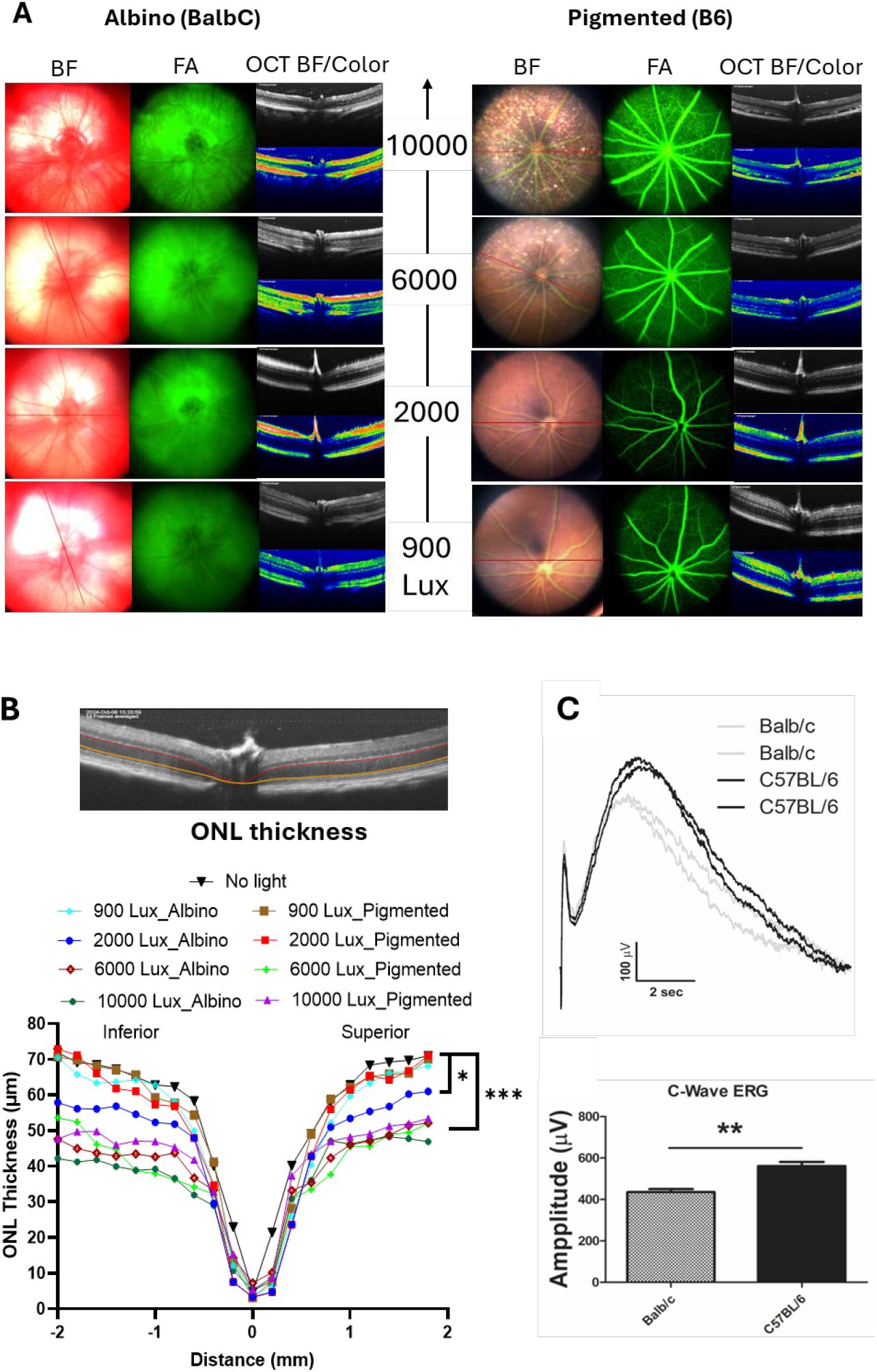
Albino (Balb/c) mice are more susceptible to blue light–induced retinal damage. (a) Schematic of the blue light exposure protocol (460 nm, 900–10,000 lux, 3 h/day for 3 consecutive days). Representative bright-field (BF) fundus images, fluorescein angiography (FA), and optical coherence tomography (OCT) acquired on day 5 after the final exposure demonstrate markedly greater photodamage in albino (Balb/c) mice compared with pigmented (B6) controls. Discrete white fundus lesions, indicative of outer retinal injury, are evident in albino mice at exposure intensities as low as 2000 lux, whereas pigmented mice show minimal changes at comparable intensities. (b) Quantification of outer nuclear layer (ONL) thickness from OCT images shows significant ONL thinning in albino mice beginning at 2000 lux, with progressively increased damage at higher light intensities (4000–10,000 lux). Pigmented mice display preserved ONL structure at lower intensities and only mild thinning at higher irradiance. (c) Quantification of RPE c-wave amplitude comparing pigmented and albino mouse eyes. ERG recordings reveal a significantly reduced c-wave amplitude in albino mice compared with pigmented controls. Data are presented from n = 5 mice per group.

### Ex vivo culture and characterization of mouse primary RPE cells

Primary RPE cells were successfully isolated and cultured from both pigmented (B6) and albino (Balb/c) mouse eyes. After two weeks in culture, light microscopy at 10× and 40× magnification showed that B6-derived RPE cells displayed the expected polygonal epithelial morphology with abundant intracellular pigmentation (Fig. 2). In contrast, Balb/c RPE cells exhibited a complete absence of pigment granules, consistent with the albino phenotype. When RPE cells were passaged to new culture plates, B6-derived cells progressively lost pigmentation, generating “de-pigmented” RPE cultures. Despite these differences in melanin content, all cultures pigmented B6, de-pigmented B6, and albino Balb/c maintained a typical cobblestone morphology with intact cell– cell junctions.

**Figure 2.**
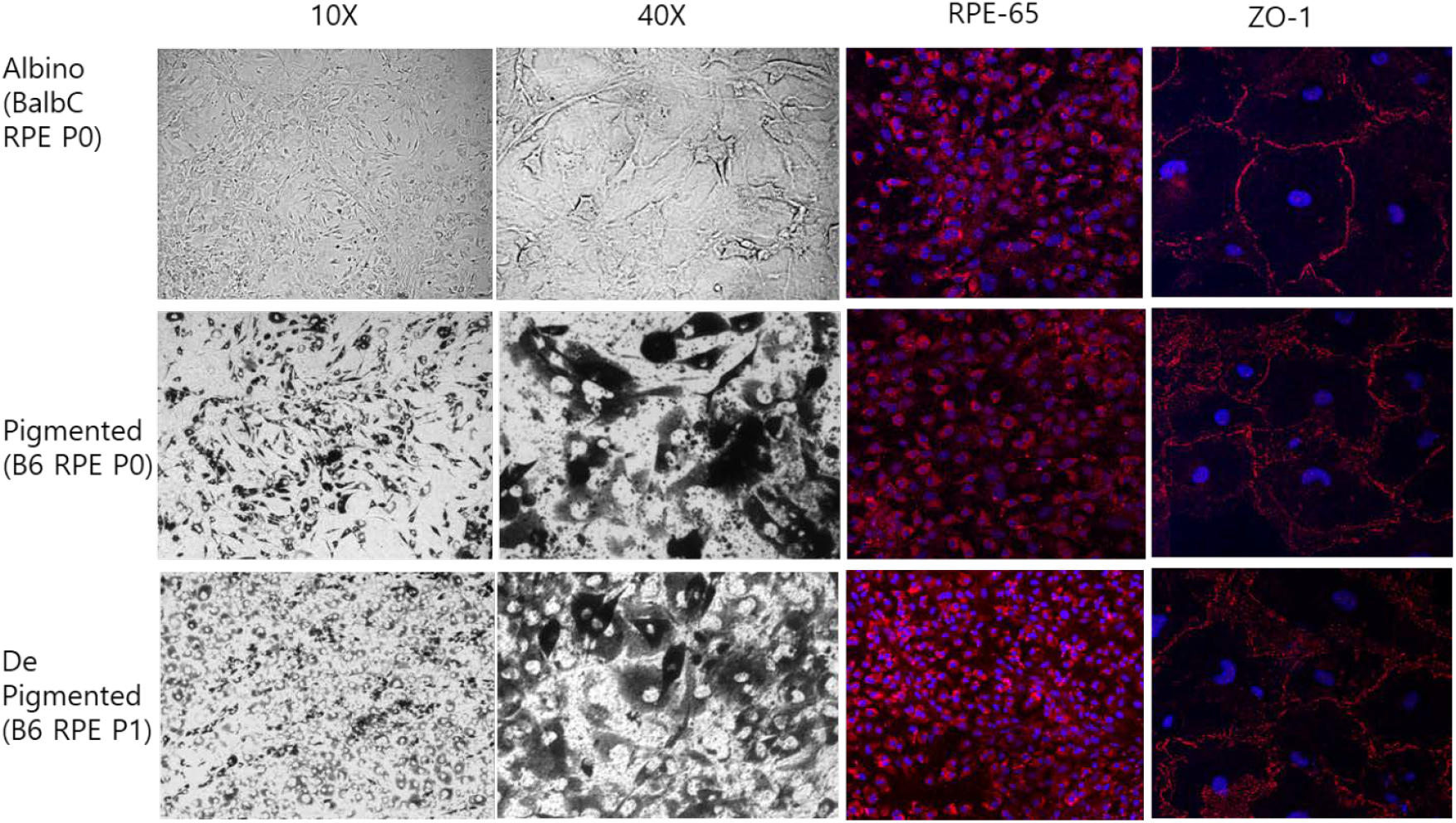
Ex vivo culture and characterization of primary RPE cells. Light microscopy images at 10× and 40× magnification show the morphology of primary RPE cultures derived from albino (Balb/c) and pigmented (B6) mice). Confocal imaging reveals expression of RPE65 and the tight junction protein ZO-1, confirming epithelial integrity

To verify RPE identity and epithelial integrity, cultures were immunoassayed for key RPE markers. Confocal imaging demonstrated strong expression of RPE65, a visual cycle protein characteristic of mature RPE, across all groups. Similarly, the tight junction protein ZO-1 formed continuous circumferential junctional belts at the apical borders, indicating preserved barrier function and epithelial polarity. Importantly, the intensity and localization of both RPE65 and ZO-1 were comparable among pigmented, de-pigmented, and albino RPE cultures, demonstrating that pigment loss during passaging or the inherent absence of pigment in albino RPE does not alter fundamental RPE cellular characteristics.

### Mitochondrial morphology and copy number differ between pigmented and albino RPE

To determine whether pigmentation status influence’s mitochondrial structure and abundance in the RPE, we examined freshly isolated RPE tissue and ex vivo primary RPE cultures derived from pigmented (B6) and albino (Balb/c) mice. Ultrastructural analysis by transmission electron microscopy (EM) revealed marked differences in mitochondrial morphology and distribution among groups (Fig. 3). Pigmented B6 RPE at passage 0 (P0) contained larger, elongated mitochondria distributed throughout the cytoplasm with well-defined cristae. In contrast, albino Balb/c RPE (P0) displayed smaller mitochondria that were more numerous and densely clustered. Sub-culturing of pigmented B6 RPE to passage 1 (P1) produced “de-pigmented” cells, which exhibited mitochondrial features more similar to pigmented RPE than to albino cells.

**Figure 3.**
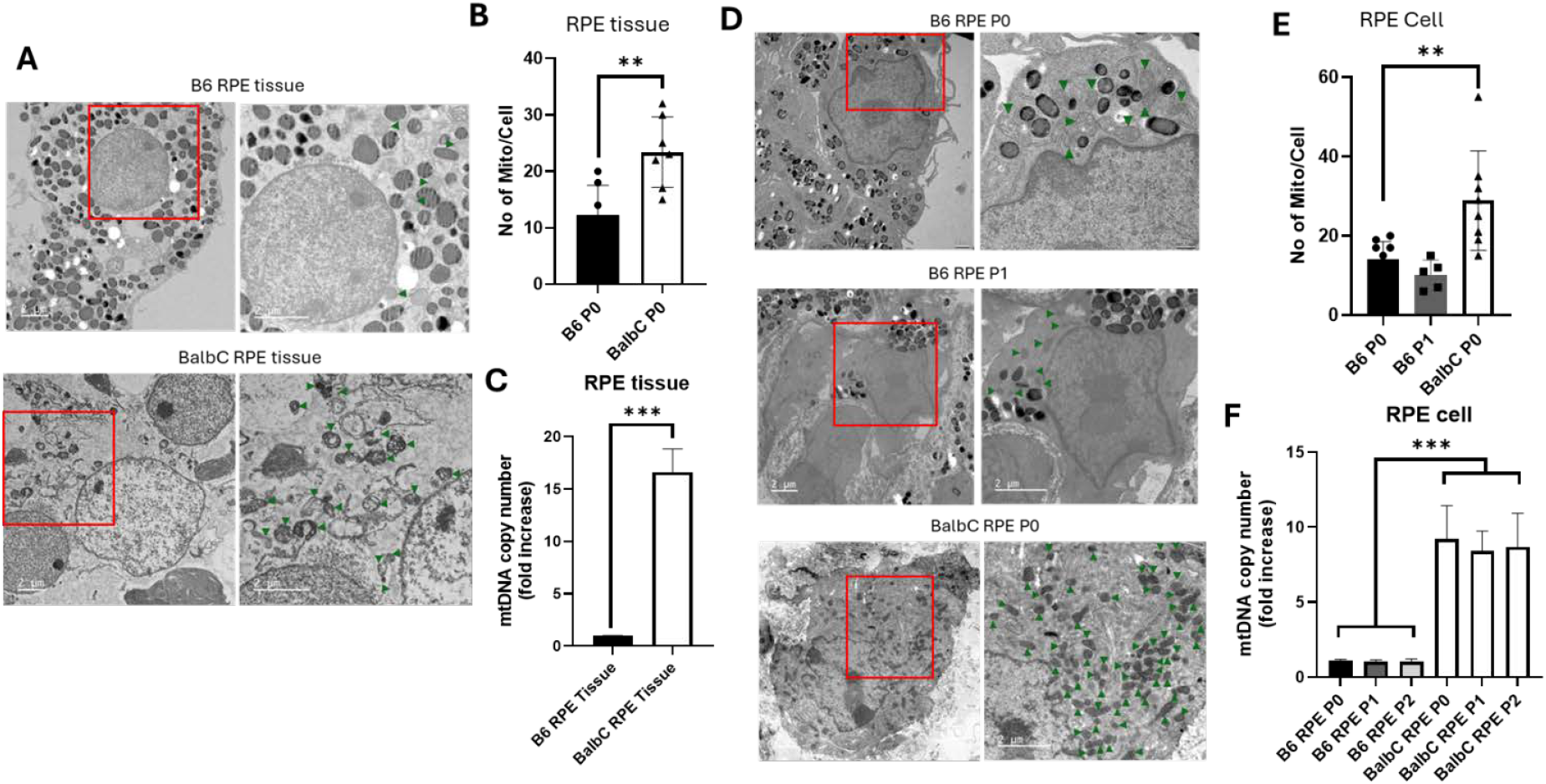
Ex vivo primary RPE cell mitochondria differ in size and number between albino and pigmented eyes. (a, d) Representative transmission electron microscopy (TEM) cross-sectional images demonstrate distinct mitochondrial distribution and morphology in pigmented (B6) and albino (Balb/c) RPE tissue, as well as in pigmented (B6 RPE P0), de-pigmented (B6 RPE P1), and albino (Balb/c RPE P0) primary RPE cells. Notable differences in mitochondrial number and morphology are observed, particularly in albino and de-pigmented RPE compared with pigmented controls (scale bar: 2 μm). (b, e) Quantitative analyses revealed that mitochondria in albino RPE tissues and cells were generally smaller and more numerous compared to those observed in pigmented and depigmented RPE. (c) Quantitative RT-PCR analysis of mitochondrial DNA (mtDNA) copy number from RPE tissue reveals significantly higher mtDNA levels in albino RPE compared with pigmented RPE. (f) In contrast, mtDNA copy number analysis from cultured RPE cell DNA shows that sub-culturing from passage 0 (P0) to passages 1 (P1) and 2 (P2) does not significantly alter mtDNA levels in either pigmented or albino RPE cells. All data are presented from groups with n = 5–7 samples per group.

Quantitative analysis confirmed that albino RPE contained significantly more mitochondria per cell than pigmented B6 RPE, both in freshly isolated tissue (Fig. 3A, B) and in primary cultures (Fig. 3D, E). These findings indicate an inherent difference in mitochondrial abundance between pigmented and albino RPE.

To assess whether mitochondrial biogenesis contributed to these differences, mitochondrial DNA (mtDNA) copy number was measured by real-time PCR using *mtCOXII* normalized to nuclear 18S rRNA. MtDNA quantification showed that mtDNA copy number was significantly higher in albino RPE compared with pigmented RPE, consistent with the increased mitochondrial abundance observed by EM. However, mtDNA levels remained stable across passages (P0–P2) in both pigmented and albino RPE (Fig. 3F), indicating that culture-induced depigmentation does not alter mtDNA content.

### Difference of cell cycle in pigmented compared to Albino RPE cell

To determine whether pigmentation status affects cell cycle dynamics in the RPE, primary RPE cells derived from pigmented and albino mice were stained with propidium iodide and analyzed by flow cytometry (Fig. 4). Cell cycle profiles were quantified and the percentage of cells in G1, S, and G2/M phases was compared between groups (Fig. 4A). Analysis of DNA content revealed no significant difference in the proportion of cells in the G1 or S phases between pigmented and albino RPE cultures (Fig. 4B). Both cell types displayed comparable G1 arrest and DNA synthesis activity, indicating similar baseline proliferative status. However, a marked increase in the G2/M population was observed in albino RPE cells compared with pigmented B6 RPE. This elevation in G2/M phase fraction suggests that albino RPE cells may exhibit delayed progression through mitosis or enhanced accumulation in G2/M. These findings indicate that although pigmentation status does not influence entry into the cell cycle (G1 or S phase), albino RPE cells show a distinct shift toward G2/M accumulation, pointing to potential differences in cell cycle regulation, checkpoint activation, or responses to intracellular stress.

**Figure 4.**
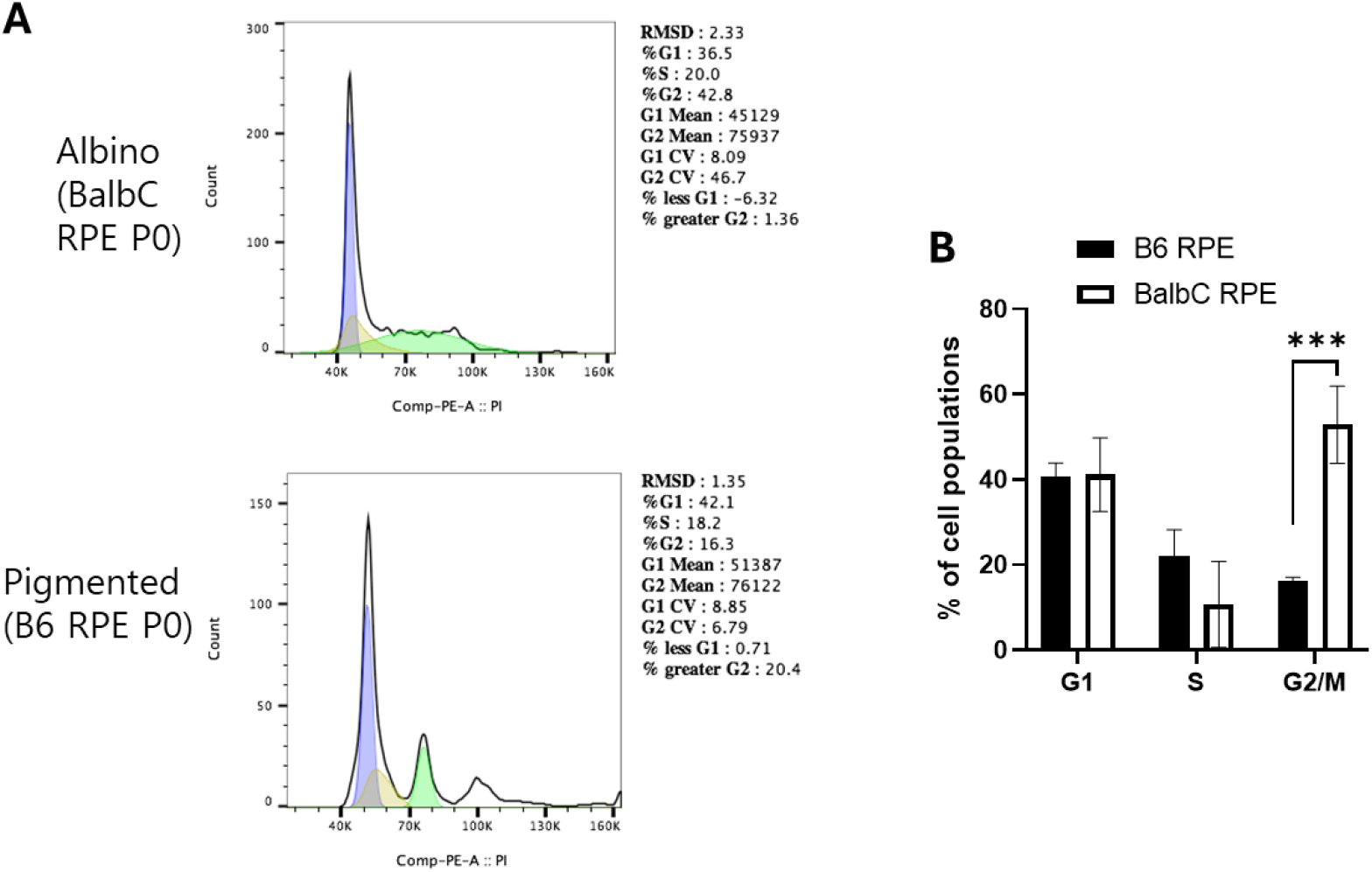
Cell-cycle differences between pigmented and albino primary RPE cells. Primary RPE cells isolated from pigmented (B6) and albino (Balb/c) mice were stained with propidium iodide and analyzed by flow cytometry to assess cell-cycle distribution. (a) Representative DNA-content histograms and (b) quantitative analysis of the percentage of cells in G1, S, and G2/M phases, determined using FlowJo. Comparative analysis reveals no significant differences in the proportion of cells in the G1 or S phases between pigmented and albino RPE cultures. In contrast, albino RPE cells exhibit a significantly increased G2/M population compared with pigmented RPE, indicating altered cell-cycle progression or accumulation at the G2/M transition. Data is presented from three independent experiments.

### Mitochondrial fission and fusion gene expression differ between pigmented and albino RPE

To determine whether pigmentation status influences mitochondrial dynamics, we examined the expression of key mitochondrial fission and fusion regulators in ex vivo primary RPE cells from pigmented (B6), de-pigmented (B6 P1/P2), albino (Balb/c), and re-pigmented (Balb/c + melanin) cultures under basal conditions and following blue-light exposure. Relative mRNA expression levels of the mitochondrial fusion genes Mfn1, Mfn2, and OPA1, and the fission gene Drp1, were quantified by qRT-PCR and normalized to control levels (Fig. 5, Supplementary Figure 1). Expanded quantitative expression data are presented in Supplementary Fig. 1. Albino RPE exhibited a significant upregulation of Drp1, indicating enhanced mitochondrial fission compared with pigmented B6 RPE. Conversely, the expression of mitochondrial fusion genes Mfn1, Mfn2, and OPA1 were markedly reduced in albino RPE. These transcriptional changes align with the ultrastructural findings of smaller, more numerous mitochondria in albino cells observed by EM (Fig. 3). De-pigmented B6 RPE (P1 and P2) showed intermediate expression patterns, including partial reductions in fusion genes and modest elevations in Drp1, suggesting that pigment loss progressively shifts mitochondrial dynamics toward a more fragmented mitochondrial network.

**Figure 5.**
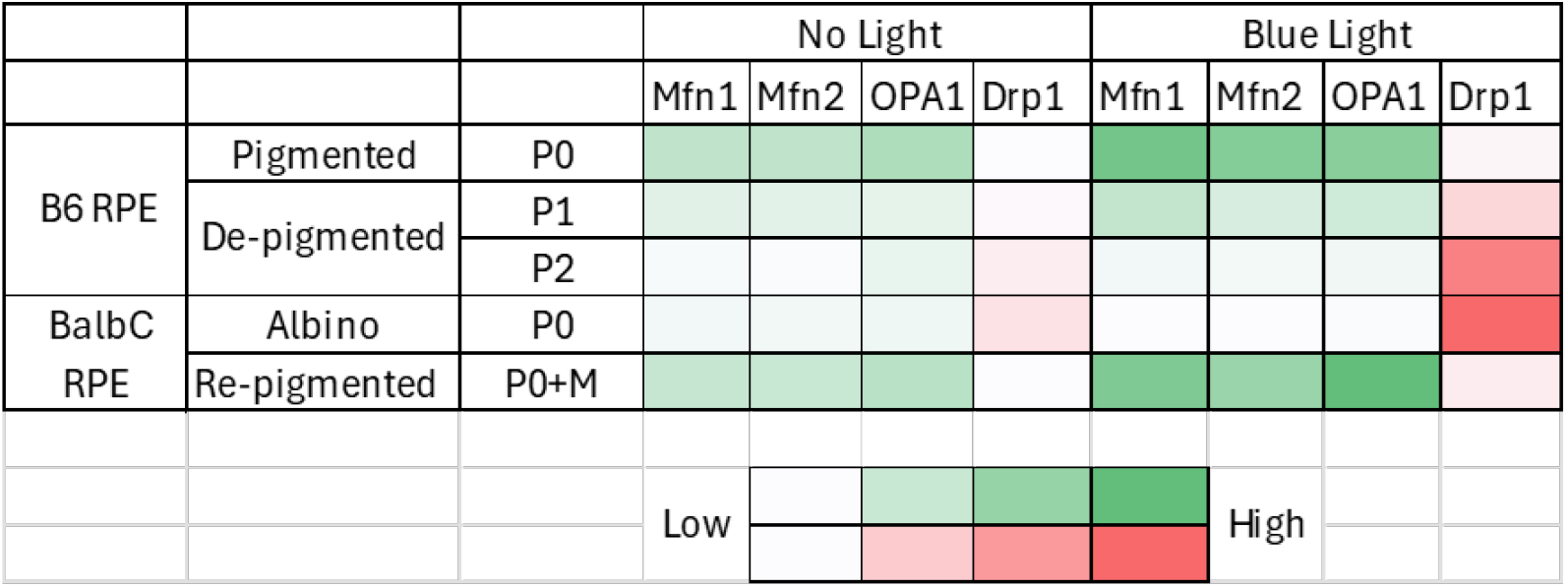
Ex-vivo primary RPE cell mitochondria is different gene expression between albino and pigmented eyes. Relative mRNA expression levels of mitochondrial dynamics regulators—Mfn1, Mfn2, OPA1, and Drp1—were assessed in pigmented (B6 RPE P0), de-pigmented (B6 RPE P1, P2), albino (Balb/c RPE P0), and re-pigmented (Balb/c RPE P0 + Melanin) RPE cells in presence or absence of blue light. Data are normalized to control levels and presented as mean ± SEM from three independent experiments. Data is presented from three independent experiments.

Re-pigmentation of albino RPE through exogenous melanin supplementation partially restored the expression balance, increasing fusion gene levels and reducing Drp1 expression. This reversal supports a functional relationship between intracellular melanin content and mitochondrial dynamics regulation.

Exposure to blue light further amplified these differences: albino and de-pigmented RPE displayed stronger induction of Drp1 and greater suppression of fusion transcripts, whereas pigmented RPE maintained relatively stable gene expression profiles. These observations are consistent with prior reports linking mitochondrial fragmentation to heightened cellular stress and increased vulnerability in RPE cells[47].

Together, these findings demonstrate that pigmentation status is a critical determinant of mitochondrial fission–fusion balance in RPE cells, with albino and de-pigmented RPE favoring a fragmented mitochondrial phenotype, and pigmented RPE maintaining a more fusion-dominant state.

### Impaired mitochondrial respiration and ATP production in albino RPE

To assess whether pigmentation status influences mitochondrial bioenergetic function in the RPE, OCR and mitochondrial ATP production were measured in RPE tissue and primary RPE cultures using the Seahorse ATP Real-Time Rate Assay (Fig. 6). Pigmented B6 RPE tissue showed significantly higher basal OCR compared with albino Balb/c RPE (Fig. 6 A, B). This increase in oxygen consumption correlated with a marked elevation in mitochondrial ATP production, indicating more efficient oxidative phosphorylation in pigmented RPE (Fig. 6 C. In contrast, albino RPE tissue exhibited reduced mitochondrial respiration and lower ATP output, consistent with compromised mitochondrial function. Similar pattern was observed in ex vivo primary RPE cultures. Pigmented B6 RPE at passage 1 (P1) displayed substantially elevated OCR (Fig. 6 D, E) and correspondingly higher levels of mitochondrial ATP generation relative to de-pigmented B6 RPE (P2 and P3) and albino Balb/c RPE (P1) (Fig. 6F). Both de-pigmented and albino RPE demonstrated diminished mitochondrial respiration, indicating that loss of pigmentation whether inherent or acquired negatively affects cellular bioenergetic capacity.

**Figure 6.**
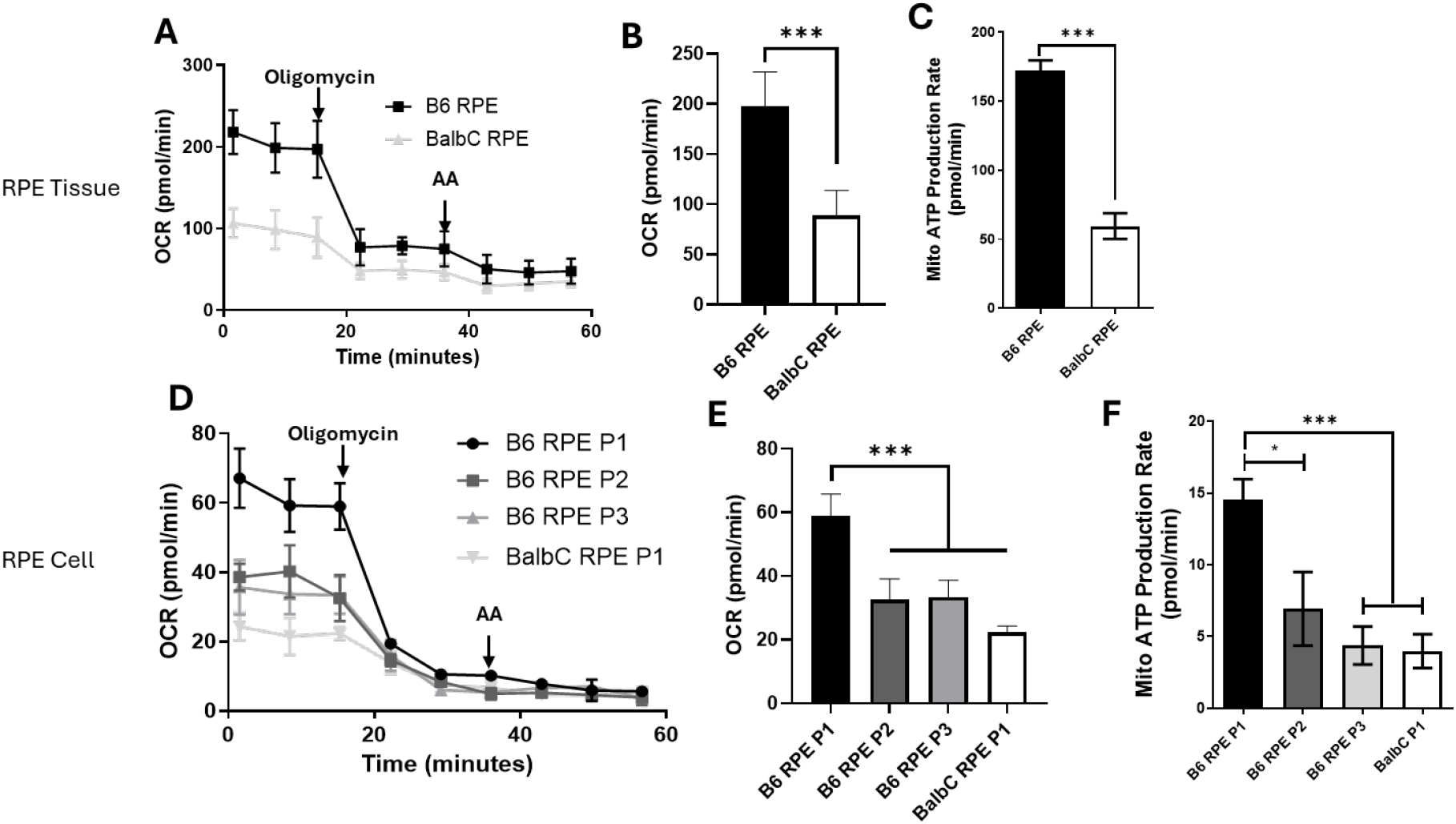
Impaired mitochondrial respiration and ATP production in Albino RPE. (a) OCR was measured using the Seahorse ATP Real-Time Rate Assay in RPE tissues. (b–c) Quantitative analyses demonstrated that pigmented (B6) RPE tissue exhibited significantly higher OCR (b) and correspondingly greater mitochondrial ATP production (c) compared to albino (Balb/c) RPE tissue. (d) OCR was similarly assessed in primary RPE cells using the Seahorse ATP Real-Time Rate Assay. (e–f) Quantitative analyses showed that pigmented primary RPE cells (B6 RPE P1) exhibited significantly increased OCR (e) and enhanced mitochondrial ATP production (f) relative to depigmented (B6 RPE P2 and P3) and albino (Balb/c RPE P1) cells. Data are presented from three independent experiments.

### Mitochondrial respiration in pigmented and re-pigmented RPE depends on MPC-mediated pyruvate import

To determine whether pigmentation influences metabolic substrate utilization in the mitochondrial respiratory chain, we measured OCR in pigmented (B6 RPE P1), de-pigmented (B6 RPE P2), albino (Balb/c RPE P1), and re-pigmented (Balb/c RPE P1 + melanin) primary RPE cells following inhibition of key metabolic pathways. Under basal conditions, OCR was highest in pigmented and re-pigmented RPE, whereas both de-pigmented and albino RPE showed markedly reduced mitochondrial respiration (Fig. 7 A), consistent with impaired bioenergetic performance.

**Figure 7.**
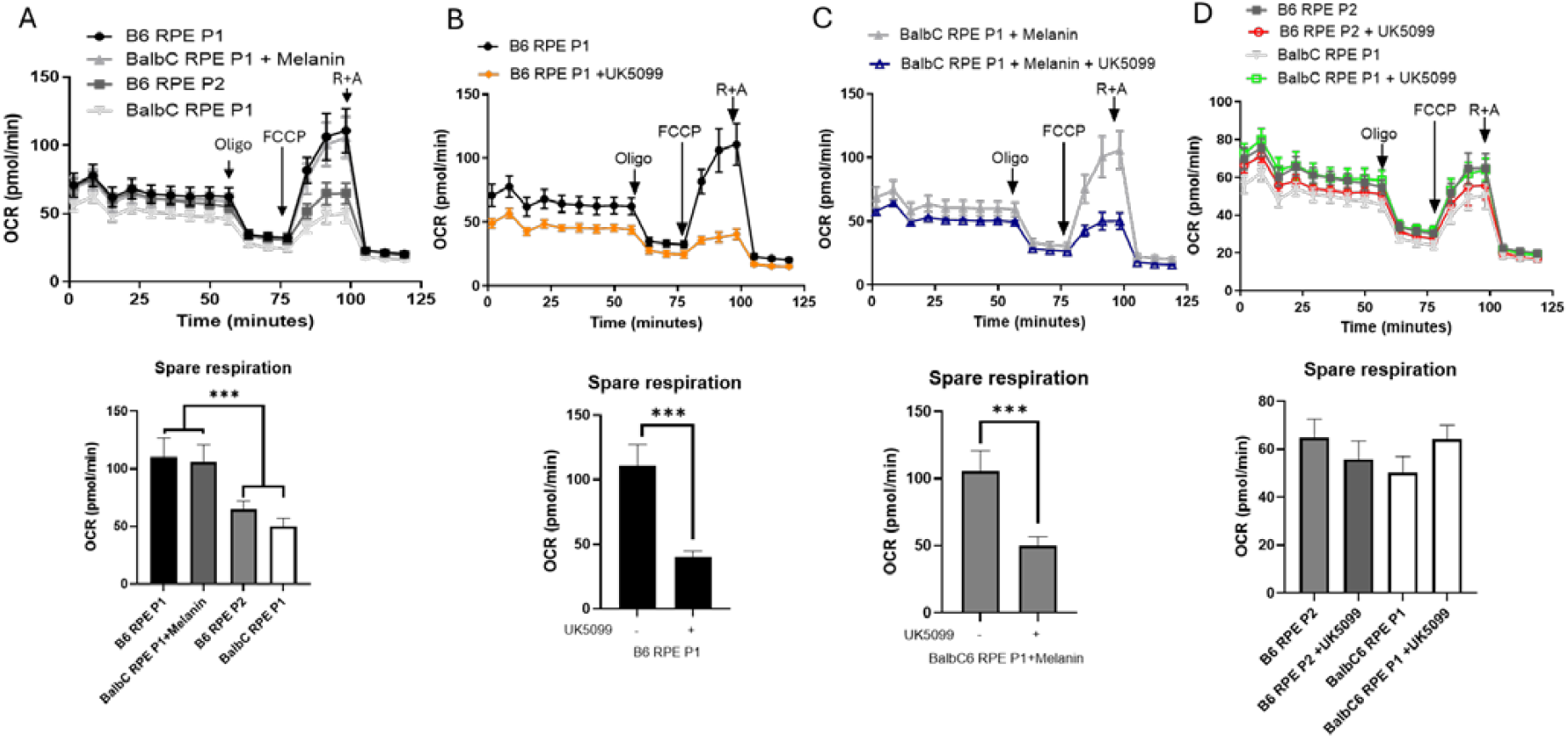
Mitochondrial respiration in pigmented and re-pigmented RPE cells depends on MPC-mediated pyruvate import. (a) Basal OCR measurements reveal significantly higher mitochondrial respiration in pigmented (B6 RPE P1) and re-pigmented (Balb/c RPE P1 + melanin) RPE cells compared with de-pigmented (B6 RPE P2) and albino (Balb/c RPE P1) RPE cells. (b, c) Inhibition of the mitochondrial pyruvate carrier (MPC) with UK5099 causes a pronounced reduction in OCR in pigmented (b) and re-pigmented (c) RPE cells, indicating a strong dependence on MPC-mediated pyruvate import to sustain mitochondrial respiration. (d) In contrast, UK5099 has minimal effect on OCR in de-pigmented and albino RPE cells, demonstrating reduced reliance on pyruvate-driven oxidative phosphorylation in the absence of melanin. Data is presented from three independent experiments.

Inhibition of the mitochondrial pyruvate carrier (MPC) with UK5099 caused a pronounced reduction in OCR in pigmented (Fig. 7 B) and re-pigmented (Fig. 7 C) RPE cells, indicating that these cells rely heavily on MPC-mediated pyruvate import to fuel mitochondrial respiration. In contrast, UK5099 had minimal effect on OCR in de-pigmented and albino RPE (Fig. 7 D), demonstrating reduced dependence on pyruvate-driven oxidative phosphorylation in cells lacking melanin. These findings suggest a potential functional link between melanin content and mitochondrial pyruvate metabolism, implicating melanin-associated energy transfer into the TCA cycle via the MPC pathway.

In contrast to MPC inhibition, Etomoxir (CPT1 inhibitor, fatty acid oxidation block) (Supplementary Fig. 2), and BPTES (glutaminase inhibitor, glutamine oxidation block) (Supplementary Fig. 3) produced similar OCR responses across pigmented, albino, de-pigmented, and re-pigmented RPE, indicating that β-oxidation and glutaminolysis do not differ significantly with pigmentation status.

Together, these findings demonstrate that pigmentation specifically enhances reliance on pyruvate-dependent mitochondrial respiration, while other metabolic pathways contribute similarly regardless of pigment content.

### Albino RPE exhibit heightened inflammatory cytokine secretion in response to blue-light exposure

To determine whether pigmentation status influences inflammatory signaling in the RPE, we quantified secretion of IL-6, TNF-α, IFNβ1, and IL-1β from pigmented (B6 RPE P0), de-pigmented (B6 RPE P1/P2), albino (Balb/c RPE P0), and re-pigmented (Balb/c RPE P0 + melanin) primary RPE cultures under basal conditions and following blue-light exposure (Fig. 8; Supplementary Fig. 4). Cytokine secretion levels were normalized to untreated controls.

**Figure 8.**
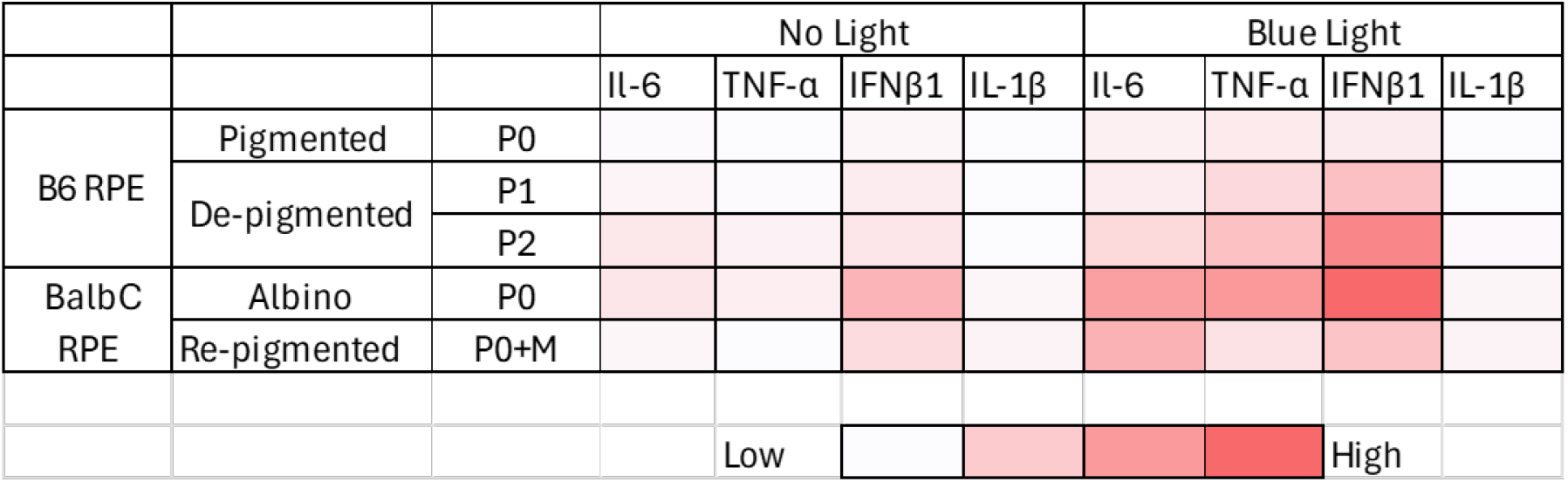
Albino and pigmented RPE cells differentially secrete inflammatory cytokines in *ex vivo* culture. Inflammatory cytokine (IL-6, TNF-α, IFNβ1 and IL-1 β) secreted by pigmented (B6 RPE P0), de-pigmented (B6 RPE P1, P2), albino (Balb/c RPE P0), and re-pigmented (Balb/c RPE P0 + Melanin) RPE cells in presence or absence of blue light. Data are normalized to control levels and presented as mean ± SEM from three independent experiments. Data is presented from three independent experiments.

Albino RPE secreted significantly higher levels of IFNβ1 compared with pigmented, de-pigmented, and re-pigmented RPE. Exposure to blue light further exacerbated IFNβ1 release specifically in albino RPE, whereas cytokine levels in pigmented and re-pigmented cells remained comparatively stable. This pronounced induction of IFNβ1 suggests increased sensitivity of albino RPE to photo stress.

Inflammatory cytokines IL-6 and TNF-α showed a similar secretion pattern. Basal levels were elevated in albino RPE relative to pigmented, de-pigmented, and re-pigmented cultures, and blue-light stimulation markedly intensified IL-6 and TNF-α secretion in albino RPE. De-pigmented B6 RPE showed a moderate increase, whereas pigmented and re-pigmented cells displayed minimal changes, indicating a protective role of melanin in suppressing inflammatory activation.

In contrast, IL-1β secretion demonstrated minimal variation between the different RPE types under both basal and blue-light conditions, suggesting that this cytokine is less responsive to pigmentation-dependent modulation.

Together, these findings indicate that albino RPE are intrinsically more prone to pro-inflammatory cytokine release, and that blue-light exposure further amplifies this inflammatory response. The reduced cytokine production in re-pigmented RPE supports a functional role for melanin in dampening photo stress-induced inflammation.

## Discussion

This study identifies melanin as a pivotal determinant of mitochondrial structure, metabolism, and inflammatory tone in the RPE, providing a mechanistic explanation for the heightened susceptibility of albino retinas to blue light–induced injury. By integrating *in vivo* retinal damage paradigms with structural, metabolic, transcriptional, and inflammatory analyses, we demonstrate that pigmentation status governs a coordinated mitochondrial program essential for retinal resilience.

Consistent with prior reports, albino Balb/c mice displayed dramatically enhanced phototoxic responses to blue light compared with pigmented C57BL/6J mice[23, 27]. Structural degeneration manifested as ONL thinning, and discrete fundus lesions at light intensities that were well tolerated by pigmented animals. These findings reinforce the protective role of melanin against photic stress but also suggest the involvement of downstream cellular mechanisms beyond optical absorption alone.

EM revealed striking differences in mitochondrial morphology between pigmented and albino RPE. Albino RPE contained significantly more mitochondria per cell, yet these organelles were smaller, fragmented, and clustered hallmarks of excessive mitochondrial fission. Elevated mtDNA copy number further supports compensatory mitochondrial biogenesis in response to inefficient energy production, a phenomenon observed in other degenerative contexts[48]. Importantly, depigmentation induced by RPE passaging partially recapitulated these phenotypes, indicating that pigment loss itself, not genetic background alone, contributes to mitochondrial remodeling.

Transcriptional profiling directly linked mitochondrial morphology to altered expression of fission–fusion regulators. Albino RPE exhibited strong upregulation of *Drp1* accompanied by suppression of *Mfn1, Mfn2*, and *OPA1*, favoring a fragmented mitochondrial network. Excessive Drp1-mediated fission has been implicated in RPE degeneration, inflammasome activation, and bioenergetic failure[49]. The partial normalization of these genes following melanin re-pigmentation provides compelling evidence that melanin actively stabilizes mitochondrial dynamics rather than serving as a passive pigment.

A major functional consequence of pigmentation-dependent mitochondrial reprogramming was revealed by ATP real-time rate analysis. Pigmented and re-pigmented RPE relied heavily on MPC-mediated pyruvate import to sustain oxidative phosphorylation, whereas albino and depigmented RPE displayed blunted responses to MPC inhibition, indicating loss of pyruvate-driven respiration. Pyruvate oxidation is a primary energy source for RPE and a key link between glycolysis and the TCA cycle[50]. Its impairment likely reduces ATP availability required for photoreceptor support and stress resistance.

Interestingly, fatty acid oxidation and glutaminolysis contributed similarly across pigmentation states, suggesting that melanin selectively enhances pyruvate utilization rather than broadly amplifying mitochondrial metabolism. This specificity implicates melanin in metabolic channeling or redox coupling that favors carbohydrate-driven respiration.

Albino RPE exhibited elevated basal and blue light-induced secretion of IFN-β1, IL-6, and TNF-α cytokines strongly associated with retinal degeneration and AMD[11, 51]. Mitochondrial fragmentation and impaired respiration are well-established drivers of type I interferon signaling and sterile inflammation[52]. The absence of differences in IL-1β suggests selective activation of mitochondrial stress-linked inflammatory pathways rather than generalized inflammasome priming. Melanin re-pigmentation significantly dampened cytokine production, highlighting its immune-modulatory function. These findings suggest that mitochondrial instability in albino RPE generates a pro-inflammatory milieu that exacerbates light-induced retinal damage.

Together, our data position melanin as an important regulator of RPE mitochondrial homeostasis, linking pigmentation to cellular bioenergetics, inflammatory restraint, and retinal protection. These results have broad implications for retinal degenerative diseases characterized by melanin loss, including AMD, albinism, and aging-related RPE dysfunction[10, 13]. Therapeutic strategies aimed at enhancing mitochondrial fusion, restoring pyruvate metabolism, or mimicking melanin’s bioenergetic effects may provide novel avenues to protect the retina from photo-oxidative injury.

## Acknowledgements

We would like to acknowledge Qing Shi from the Seahorse Core Services, Paul Risteff for TEM training and assistance with image acquisition and sample preparation, and the Histology Core Services at the University of North Carolina at Chapel Hill for histological sample preparation. We are grateful for their support and contributions to this investigation.

