## Supplementary material for "Melanin regulates mitochondrial dynamics, metabolism and inflammatory signaling to protect the retina": 05012026 Supplementary File.docx

**^a^ Department of Ophthalmology, the University of North Carolina School of Medicine, Chapel Hill. NC 27599, USA**

**^b^Division of Pharmacoengineering & Molecular Pharmaceutics, Eshelman School of Pharmacy, University of North Carolina at Chapel Hill, NC 27599, USA**

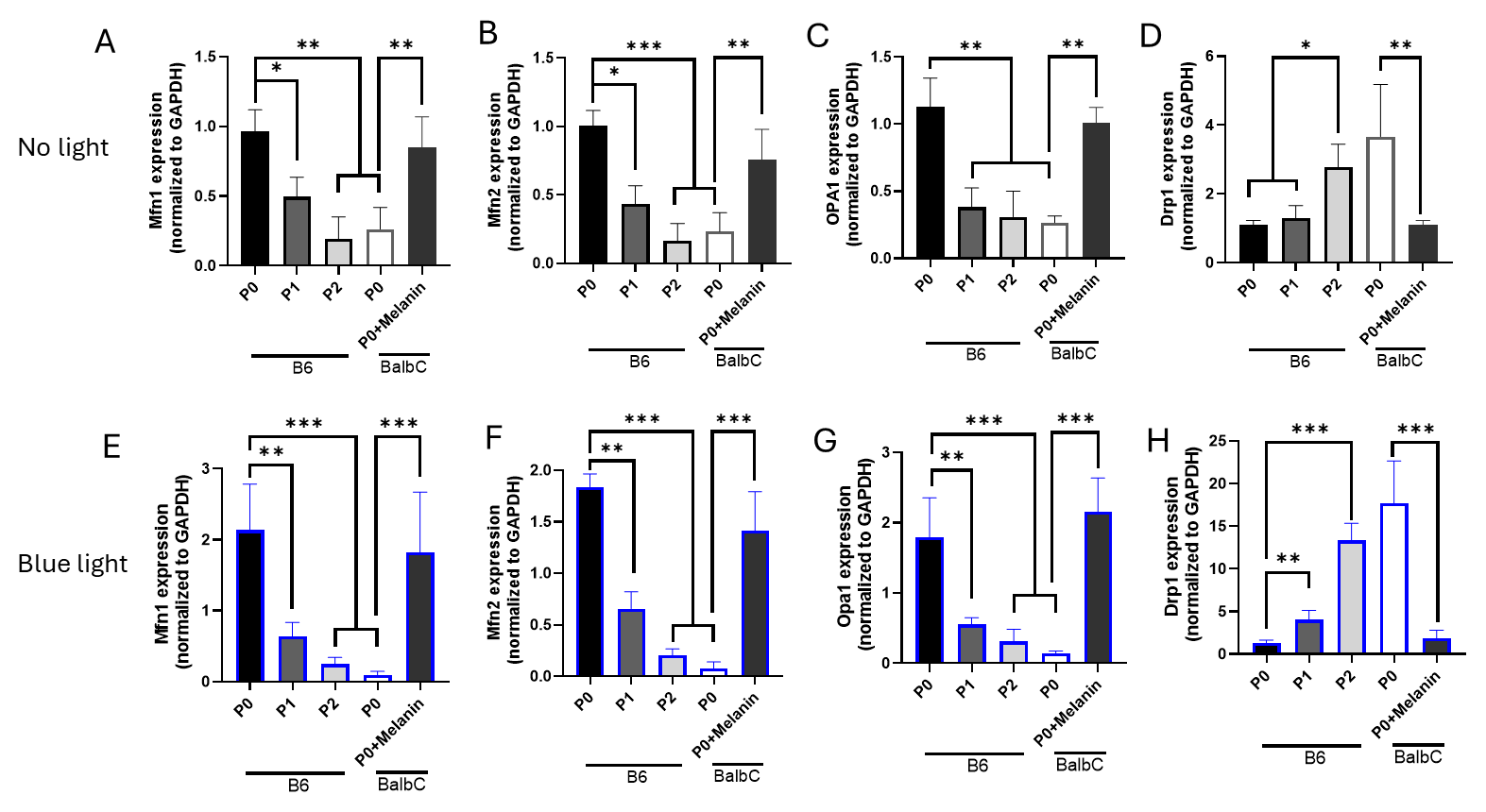

**Supplementary Figure 1. Mitochondrial fission and fusion gene expression differs between pigmented and albino primary RPE cells.**
Relative mRNA expression levels of mitochondrial dynamics regulators were analyzed in ex vivo primary RPE cells from pigmented (B6 RPE P0), de‑pigmented (B6 RPE P1, P2), albino (Balb/c RPE P0), and re‑pigmented (Balb/c RPE P0 + melanin) cultures by qRT‑PCR.
(a–d) Under basal (no‑light) conditions, expression of the mitochondrial fusion genes Mfn1 (a), Mfn2 (b), and OPA1 (c) is reduced in albino and de‑pigmented RPE cells, while the fission gene Drp1 (d) is significantly upregulated compared with pigmented and re‑pigmented RPE.
(e–h) Following blue light exposure, these differences are further accentuated, with decreased expression of Mfn1 (e), Mfn2 (f), and OPA1 (g), and a marked increase in Drp1 (h) in albino RPE cells relative to pigmented controls. Data is presented from three independent experiments.

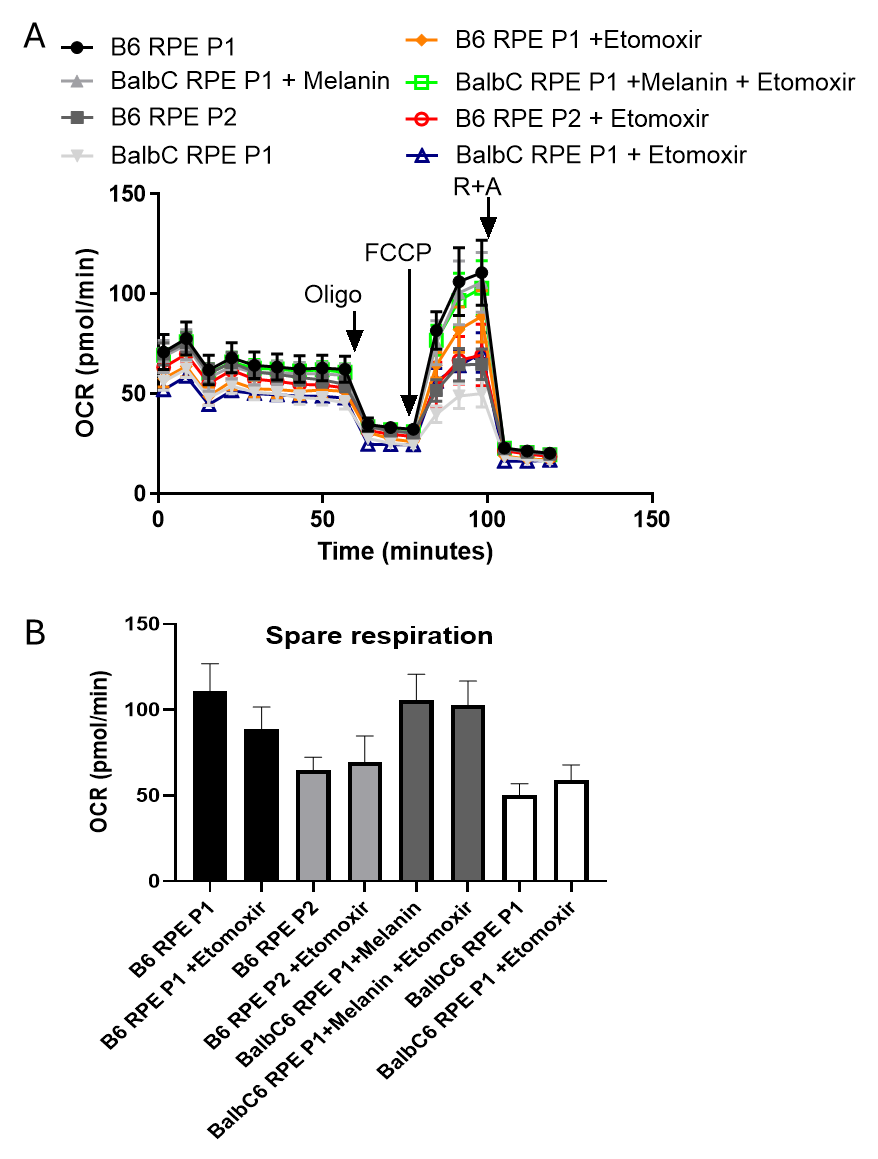

**Supplementary Figure 2. Fatty acid oxidation does not differentially contribute to mitochondrial respiration across RPE pigmentation states.**

Oxygen consumption rate (OCR) was measured in primary RPE cells from pigmented (B6 RPE P1), de‑pigmented (B6 RPE P2), albino (Balb/c RPE P1), and re‑pigmented (Balb/c RPE P1 + melanin) cultures following treatment with Etomoxir, an inhibitor of carnitine palmitoyltransferase 1 (CPT1). Data is presented from three independent experiments.

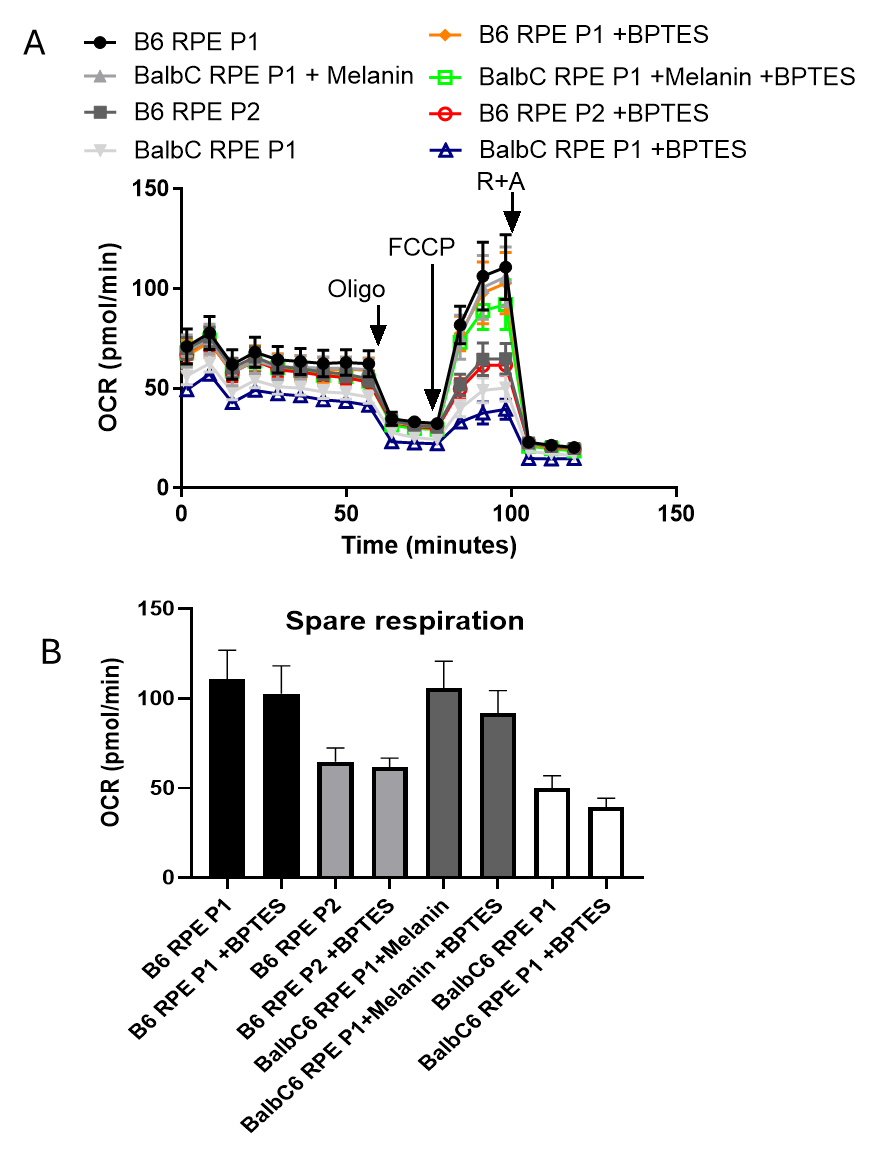

**Supplementary Figure 3. Glutamine oxidation does not differentially contribute to mitochondrial respiration across RPE pigmentation states.**

Oxygen consumption rate (OCR) was assessed in pigmented (B6 RPE P1), de‑pigmented (B6 RPE P2), albino (Balb/c RPE P1), and re‑pigmented (Balb/c RPE P1 + melanin) primary RPE cells following inhibition of glutaminase with BPTES. In contrast to MPC inhibition, BPTES elicited similar OCR responses across all pigmentation states. Data is presented from three independent experiments.

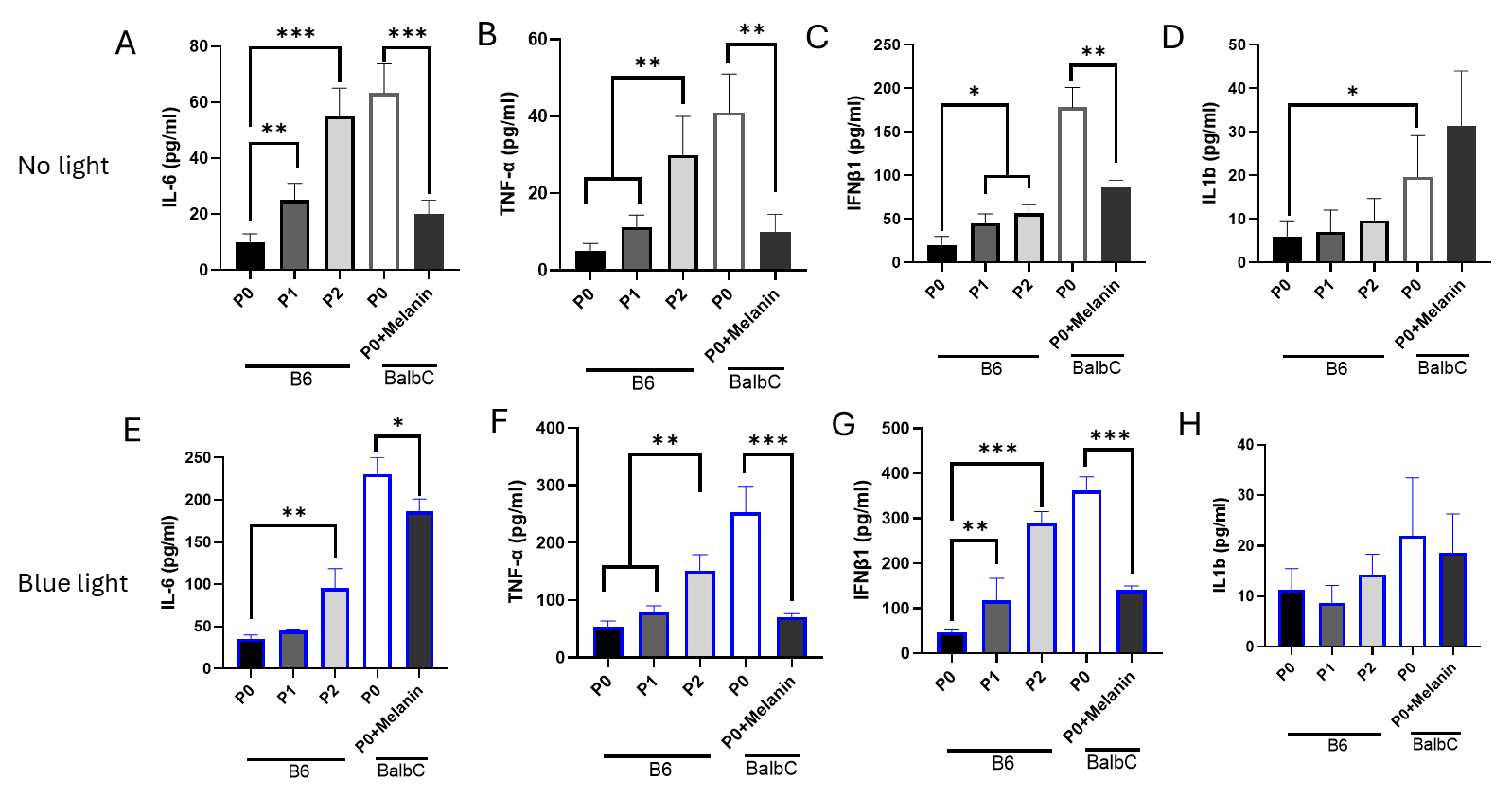

**Supplementary Figure 4. Albino and pigmented RPE cells differentially secrete inflammatory cytokines in ex‑vivo culture.**

Inflammatory cytokine secretion was quantified from primary RPE cells derived from pigmented (B6 RPE P0), de‑pigmented (B6 RPE P1, P2), albino (Balb/c RPE P0), and re‑pigmented (Balb/c RPE P0 + melanin) cultures under basal conditions or following blue light exposure.
(a–d) Under basal (no‑light) conditions, secretion levels of IL‑6 (a), TNF‑α (b), IFN‑β1 (c), and IL‑1β (d) reveal elevated IL‑6, TNF‑α, and IFN‑β1 release in albino RPE compared with pigmented and re‑pigmented controls, while IL‑1β levels remain relatively unchanged.
(e–h) Following blue light exposure, secretion of IL‑6 (e), TNF‑α (f), and IFN‑β1 (g) is markedly increased in albino RPE, indicating heightened sensitivity to photo‑stress–induced inflammation. In contrast, pigmented and re‑pigmented RPE display minimal cytokine induction, while IL‑1β secretion (h) remains largely unaffected across groups. Data is presented from three independent experiments.
